# Intracellular Delivery of Peptides and Proteins with an Engineered Membrane Translocation Domain

**DOI:** 10.64898/2026.02.13.705776

**Authors:** Prabhat Bhat, Heba Salim, Jeremy L. Ritchey, Na Li, Brendan B. Harty, Thomas Patel, Jing Zhao, Qi-En Wang, Virginia L. King, Louis Tartaglia, Jenő Gyuris, Dehua Pei

## Abstract

Antibodies and other protein therapeutics have revolutionized medicine, but their application is largely limited to extracellular targets. The lack of efficient intracellular delivery methods remains a major bottleneck. Here, we engineered a family of small (∼90 amino acids), metabolically stable membrane translocation domains (MTDs) by modifying the loop sequences of a human fibronectin type III (FN3) domain. The most potent variant, MTD4, is highly cell-permeable and can be recombinantly fused to the N- or C-terminus of any peptide or protein, serving as a versatile “plug-and-play” vehicle. We demonstrate that MTD4 fusions efficiently deliver a wide variety of functional peptides and proteins into the cytosol and nucleus of eukaryotic cells, both *in vitro* and *in vivo*. Following systemic administration, MTD4 fusion proteins exhibit broad biodistribution and homogenous tissue penetration in mice. Importantly, MTD4 is effective at low nanomolar (nM) concentrations, making it a promising platform for addressing a vast range of intracellular and previously “undruggable” targets.

## INTRODUCTION

Effective delivery of proteins into the cytosol, nucleus, or subcellular organelles (e.g., mitochondria) of mammalian cells would unlock their enormous potential for applications in protein replacement therapy,^1^ gene editing,^2^ modulation of protein-protein interactions (PPIs),^3^ and degradation of disease-causing proteins.^4^ Over the past few decades, researchers have explored a variety of approaches, including cell-penetrating peptides (CPPs),^5^ bacterial toxins,^6^ viruses,^7^ polyplexes,^8^ liposomes,^9^ and nanoparticles,^10^ to deliver proteins or their encoding DNAs/mRNAs. While some of the systems, most notably lipid nanoparticles for siRNA and mRNA,^11^ viral vectors for gene therapy and gene-editing enzymes,^12^ and bacterial toxins for anti-cancer proteins,^13^ have demonstrated efficacy in the clinic, significant challenges remain. These include the inadequate biodistribution of viral, liposomal, and nanoparticle-based systems to extrahepatic tissues, immunogenicity associated with viral vectors and bacterial toxins, and endosomal entrapment of non-viral vectors.^14-16^

An attractive approach to intracellular protein delivery is conjugation with CPPs, which are short peptides of 5–30 amino acids,^5^ because of their small sizes, biocompatibility, simplicity, and generality (i.e., the same CPP may be used to deliver many different proteins and into most cell types).

CPPs such as trans-activator of transcription (Tat),^17^ penetratin,^18^ and nonaarginine (R9)^19^ have been widely used to deliver peptides and proteins into eukaryotic cells *in vitro*. Unfortunately, the *in vivo* applications of linear CPPs have been hampered by their generally low cytosolic entry efficiency (due to endosomal entrapment) and poor metabolic stability^20^ (Figure 1). To overcome these limitations, researchers have developed structurally constrained CPPs with both improved cell entry efficiency and superior proteolytic stability. One class of structurally constrained CPPs are cyclic CPPs,^21-23^ which prove highly effective for delivering proteins and oligonucleotides into eukaryotic cells *in vivo*^24,25^ and have advanced into clinical development. However, because of their cyclic structures and often the presence of non-proteinogenic amino acids, cyclic CPPs are not genetically encodable and must be chemically synthesized and posttranslationally conjugated to a protein of interest (POI). Efficient and site-specific modification of a protein to give a single product remains a significant challenge of its own right.^26^ Another class of structurally constrained CPPs is cell-permeable miniature proteins, as exemplified by ZF5.3, which was engineered by grafting five arginine residues onto the α-helix of a zinc finger domain.^27^ ZF5.3 is genetically encodable and has been recombinantly fused to the N- or C-terminus of cargo proteins to efficiently deliver the latter into the cytosol of mammalian cells.^28,29^ A drawback of ZF5.3 is that the Zn^2+^ ion, required for folding and activity, may be lost during circulation in vivo. Additionally, it was reported that following endocytosis, ZF5.3 and the attached cargo protein must unfold in order to escape from the endosome, limiting its cargo capacity to proteins that undergo facile unfolding at physiological temperature (e.g., 37 °C).^30,31^ Finally, a third approach to generating structurally constrained CPPs involves replacing the surface-exposed loops of a POI with short CPP motifs.^32^ Grafting a CPP onto a rigid protein scaffold imposes conformational constraints on the CPP. This structural modification enhances its proteolytic stability, membrane-binding affinity, and cell-penetrating activity. While this strategy bypasses the need for post-translational modifications of the POI, it necessitates prior knowledge of the protein’s structure and may compromise the protein’s native activity or stability. Lastly, a common limitation of previous CPPs, including those with structural constraints, is their requirement of relatively high concentrations (≥1 μM) for efficacy, which restricts their application in therapeutic delivery.

**Figure 1.**
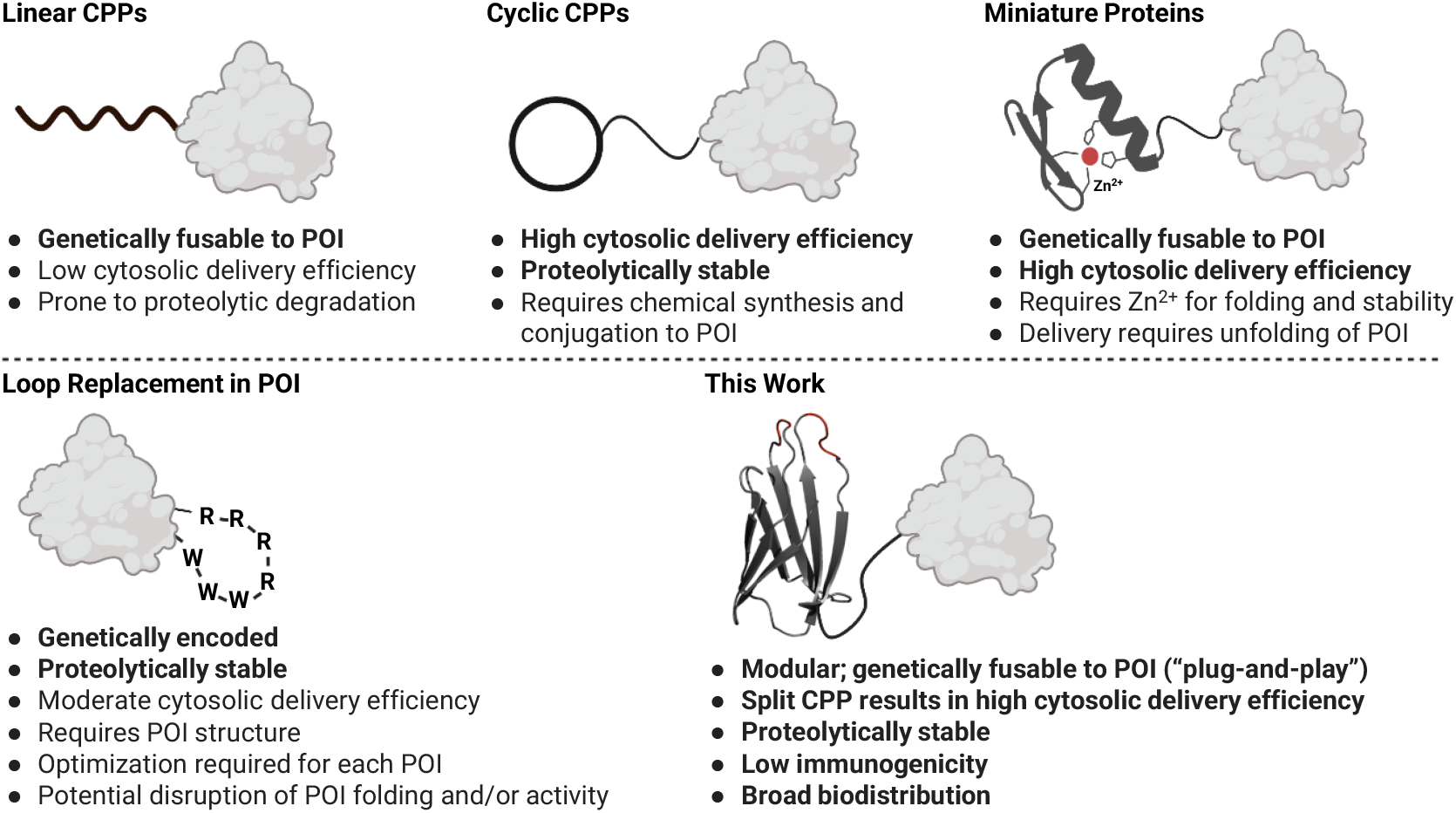
Advantages and disadvantages of different CPP-based methods for intracellular delivery of proteins.

In this study, we developed a family of small (∼90 amino acids), metabolically stable, and cell-permeable membrane translocation domains (MTDs). The most potent variant, MTD4, can be recombinantly fused to either the N- or C-terminus of any peptide or protein. This fusion effectively delivers the cargo into the cytosol of eukaryotic cells, both *in vitro* and *in vivo*. This modular, “plug-and-play” approach is highly efficient, convenient, and broadly applicable for the intracellular delivery of peptides, proteins, and potentially other biomolecules, without requiring modification of the cargo itself. Importantly, MTD4 is effective across a wide range of concentrations, delivering functional proteins at low nanomolar (nM) levels.

## RESULTS AND DISCUSSION

### Engineering Human FN3 Domain into MTDs

For our MTD scaffold, we chose the tenth fibronectin type III (FN3) domain from human fibronectin. This 94-amino-acid domain is exceptionally stable, spontaneously folds into its native conformation, and lacks cysteines or disulfide bonds, ensuring stability within the intracellular environment. It can also be efficiently produced in *Escherichia coli* and - being derived from an abundant human extracellular protein - carries minimal immunogenicity risk. These qualities have made the FN3 domain a popular scaffold for engineering monobodies, which bind target proteins with antibody-like affinity and specificity.^33^

Previous studies indicated that the BC, DE, and FG loops of the FN3 domain are highly tolerant for mutagenesis.^34^ Leveraging this and our recent discovery that amphipathic CPPs efficiently translocate cell membranes by a vesicle budding-and-collapse (VBC) mechanism,^35-37^ we designed several MTD variants. MTD1 was generated by replacing the RGDSPAS sequence in the FG loop with the CPP motif, RRRRWWW (Figure 2a, b). This amphipathic motif has previously been shown to endow membrane permeability when grafted to the surfaces of various proteins.^32^ Similarly, MTD2 resulted from substituting the AVTVR sequence in the BC loop with WWWRRRR. To assess the mutability of other loops, we replaced the tripeptide NSP of the CD loop with RRRRWWW to obtain MTD3. For MTD4, we split the CPP motif into hydrophobic (WYW) and cationic (RRRR) fragments and grafted them into the BC and FG loops, respectively, anticipating that spreading the CPP residues over two adjacent loops would increase their accessibility to the cell membrane. Finally, MTD5 was designed by swapping the RRRR and WYW motifs of MTD4. The WYW motif, being less hydrophobic than WWW, has previously been shown to aid endosomal escape of antibodies.^38^ In silico analysis suggests that in all MTDs, the CPP motifs adopt a constrained cyclic topology with their side chains exposed to the solvent. When expressed in *E. coli*, MTD2–5 yielded soluble proteins in modest to good yields, though MTD1 did not produce any soluble protein (Figure 2b).

**Figure 2.**
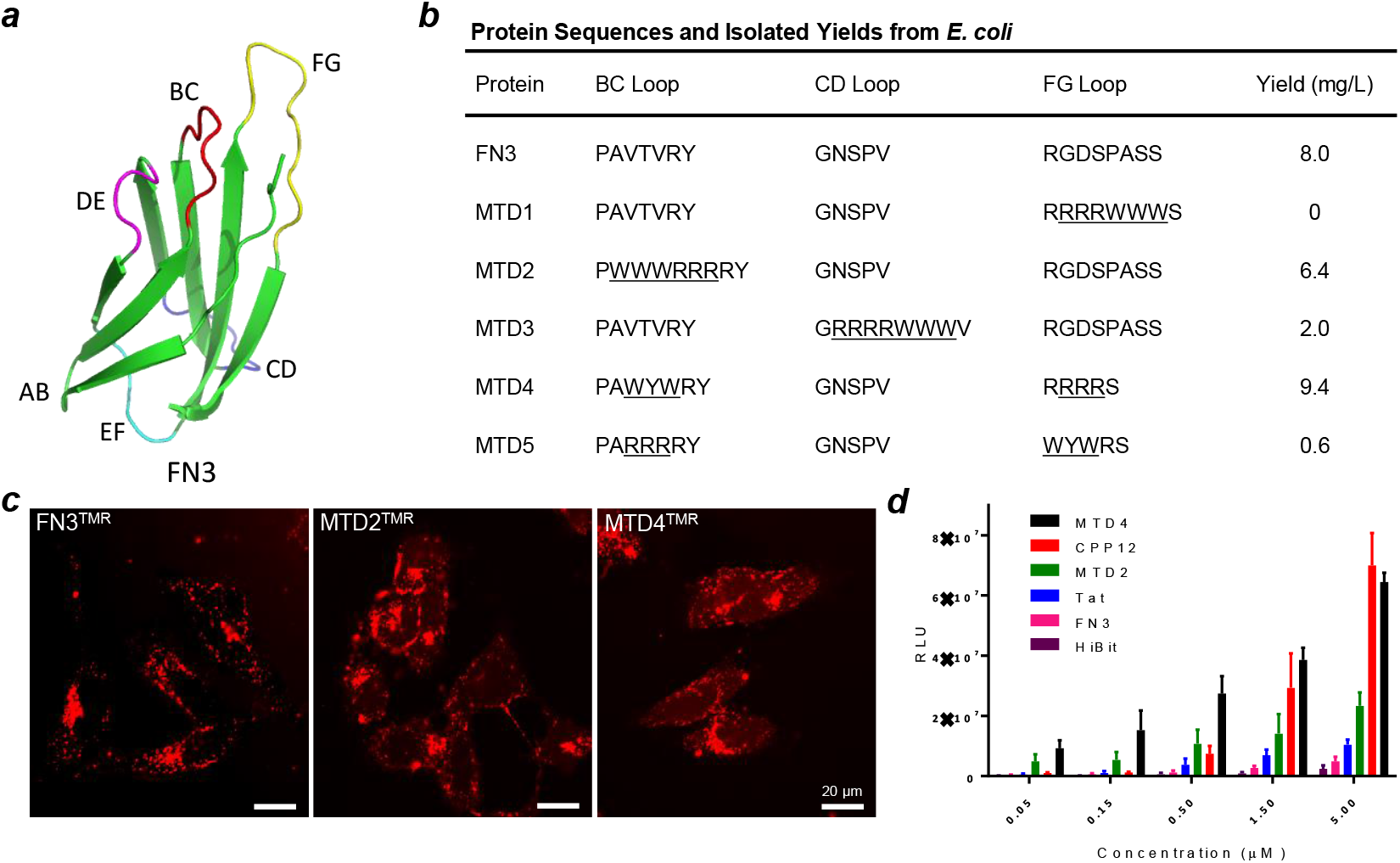
Design and characterization of MTDs. (***a***) Structure of the FN3 domain (PDB: 1ttg), with loops connecting adjacent β-strands indicated as AB, BC, CD, DE, EF, and FG. (***b***) Amino acid sequences in the BC, CD, and FG loops and isolated yields (soluble protein from *E. coli* lysate) of FN3 and MTD1–5, with the grafted CPP motifs underlined. (***c***) Live-cell confocal microscopy images of HeLa cells after incubation with 5 µM TMR-labeled proteins for 2 h. Scale bar, 20 µm. (***d***) Cytosolic delivery efficiencies of MTDs and controls as measured by the NanoLuc complementation assay. Relative luminescence units (RLU) in HEK293T cells are plotted as a function of peptide/protein concentration (n = 6).

### MTD4 Efficiently Enters the Cytosol of Eukaryotic Cells

To assess the cellular uptake and intracellular localization of the MTDs, we labeled the purified MTDs (MTD2, MTD4) and the FN3 scaffold with tetramethylrhodamine-5-maleimide (TMR) at a single cysteine introduced at their C-termini. HeLa (human cervical cancer) cells were then treated with 5 μM FN3^TMR^, MTD2^TMR^, or MTD4^TMR^ for 2 h and imaged using live-cell confocal microscopy. All three proteins showed strong intracellular fluorescence (Figure 2c). However, their localization patterns varied significantly, providing insights into their endosomal escape. FN3^TMR^ produced only punctate fluorescence, indicating that it was largely entrapped within endosomes. In contrast, MTD4^TMR^ displayed both punctate and diffuse fluorescence throughout the cell volume, including the nucleus. This diffuse pattern strongly suggests that a considerable portion of internalized MTD4^TMR^ successfully escaped the endosome and reached the cytosol and nucleus. MTD2^TMR^ showed an intermediate fluorescence pattern between FN3^TMR^ and MTD4^TMR^. Unfortunately, we couldn’t obtain sufficient labeled MTD3 or MTD5 for these studies due to protein precipitation during TMR labeling.

Next, we quantitatively measured the cytosolic delivery efficiencies of FN3, MTD2, and MTD4 using a luciferase complementation assay. ^39^ We recombinantly fused an 11-residue peptide, HiBit (corresponding to the C-terminus of NanoLuc luciferase), to the C-termini of FN3, MTD2, and MTD4 via a flexible (GGS)_3_ linker (Figure S1). For comparison, we also chemically synthesized HiBit and conjugated it to Tat, a prototypical linear CPP, and CPP12,^23^ one of the most efficient cyclic CPPs. HEK293T (human kidney) cells were transMfecTtDe5d with LgBit, the N-terminal fragment of NanoLuc, and then treated with the various HiBit-conjugated proteins and peptides. Successful delivery of HiBit into the HEK293T cell cytosol enables it to complement LgBit, forming an active luciferase that produces a measurable luminescence signal. It should be noted that conjugation of HiBit to a CPP or MTD may affect the efficiency of LgBit/HiBit complementation. This effect was estimated for each CPP or MTD by performing the complementation assay in vitro in the absence and presence of a crude HEK293T cell lysate and factored into the reported cytosolic entry efficiencies (Figure S1).

As expected, all HiBit conjugates caused dose-dependent increases in luciferase activity, though to different extents (Figure 2d). At high concentrations (e.g., 5 μM), MTD4 showed a cytosolic delivery efficiency comparable to that of CPP12 and ∼5-fold higher than that of Tat. Interestingly, as the peptide/protein concentration decreased, the amount of luminescence generated by CPP12-HiBit and Tat-HiBit dropped precipitously, whereas the declination for MTD4-HiBit was more gradual. Consequently, at low concentrations (≤ 0.15 μM), while Tat-HiBit and CPP12-HiBit produced negligible luciferase activity, MTD4 remained highly active, exhibiting a cytosolic delivery efficiency 12-, 14-, and 23-fold higher than those of CPP12, Tat, and FN3, respectively. MTD2 consistently showed approximately 3-fold lower activity than MTD4 across all tested concentrations.

### MTD4 is Thermodynamically and Proteolytically Stable

Thermal denaturation studies of MTD4 revealed a melting temperature (T_M_) of 53 ± 2 °C (Figure S2). While this T_M_ is lower than that of the native FN3 scaffold (80 ± 3 °C), MTD4 still maintains a high degree of thermodynamic stability. To evaluate its proteolytic stability, we first tested the original MTD4 construct, which included an N-terminal six-histidine tag followed by a thrombin cleavage site and a C-terminal (GGS)_3_C linker (Figure S3). This initial construct exhibited a relatively short half-life (*t*_1/2_) of ∼3 h in human serum (Figure S3). Mass spectrometric analysis revealed two primary sites of proteolysis: one at the N-terminal thrombin cleavage site and another at an R/T dipeptide located immediately N-terminal to the (GGS)_3_C linker. Removal of both proteolytic sites by mutagenesis resulted in MTD4s (“s” for “stable”), which retained the full cell-penetrating activity of MTD4 but was significantly more stable in human serum (*t*_1/2_ >24 h; Figure S3).

### MTD4 Facilitates Cytosolic Delivery of an Active Enzyme

To demonstrate the capability of MTD4 for intracellular protein delivery, we first fused the 35-kD catalytic domain of protein tyrosine phosphatase 1B (PTP1B)^40^ to its C-terminus. Since tyrosine phosphorylation primarily occurs in the cytosol and nucleus of eukaryotic cells, a reduction in the global phosphotyrosine (pY) levels of intracellular proteins serves as a direct indicator of successful PTP1B delivery into the cytosol. Western blot analysis using an anti-pY antibody revealed that MTD4-PTP1B dose-dependently reduced pY levels in HEK293T cells, achieving an EC_50_ value of ∼5 nM and almost complete dephosphorylation of all pY proteins at 500 nM (Figure 3). In contrast, treatment with PTP1B alone or a catalytically inactive mutant, MTD4-PTP1B(C215S), had no significant effect. This clearly demonstrates MTD4’s ability to deliver active enzymes to their intracellular targets.

**Figure 3.**
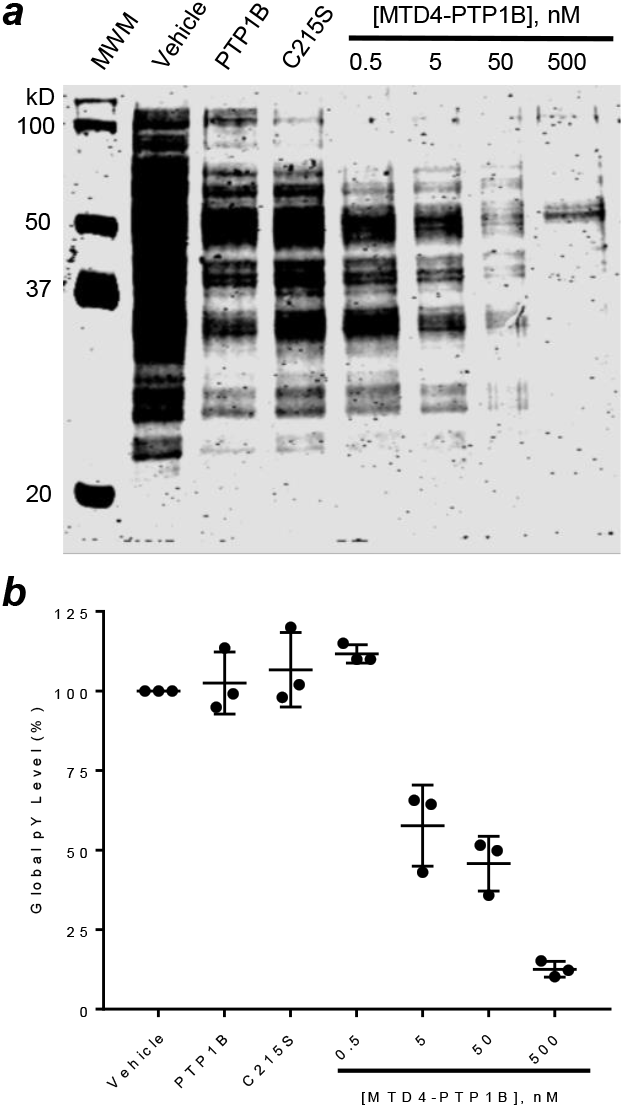
Delivery of PTP1B into the cytosol of mammalian cells. (***a***) Anti-pY Western blot of HEK293T cells after 6 h treatment with vehicle (buffer only), MTD4-PTP1B (0.5–500 nM), MTD4-PTP1B(C215S) (500 nM), or PTP1B (500 nM). MWM, molecular weight markers; C215S, MTD4-PTP1B(C215S). (***b***) Quantification of Western blot data from (***a***), normalized to pY levels of vehicle-treated cells (n = 3).

### MTD4 Delivers PPI Inhibitor of Ras Signaling

Intracellular PPIs represent a large class of exciting yet particularly challenging drug targets.^3^ To demonstrate MTD4’s utility in delivering PPI inhibitors, we engineered MTD4-RBDV, by fusing MTD4 to an optimized variant of the Ras-binding domain of C-Raf (RBDV). RBDV is a potent and selective inibitor of Ras, binding strongly to GTP-bound KRas, HRas and NRas isoforms (e.g., *K*_D_ ∼3 nM for HRas) and effectively blocking Ras’s interaction with its effector proteins.^41^ Prior work has shown that ectopic expression of RBDV in Ras-mutant cancer cells suppresses Ras signaling and induces apoptosis. ^41^

We first evaluated MTD4-RBDV’s ability to inhibit Ras-Raf interaction in live cells using a bioluminescence resonance energy transfer (BRET) assay.^42^ HEK293T cells were co-transfected with plasmids encoding KRas-luciferase and c-Raf RBD-green fluorescent protein (GFP) fusion proteins. A high BRET signal arises when KRas binds to c-Raf RBD bringing the luciferase and GFP into proximity. An inhibitor of this interaction, therefore, would reduce the BRET signal As hypothesized, MTD4-RBDV dose-dependently decreased the BRET signal in HEK293T cells expressing either KRas(G12V) or KRas(G12D), with IC_50_ values ranging from 5–10 μM (Figure 4a, b). Consistent with its ability to inhibit Ras-Raf interaction, MTD4-RBDV also dose-dependently reduced the viability of all tested Ras-mutant cancer cell lines including H358 (lung cancer), A549 (lung cancer), Mia PaCa-2 (pancreatic ductal adenocarcinoma), H1915 (lung carcinoma), H1299 (lung carcinoma), and SW480 (colorectal cancer), with IC_50_ values between 1–5 μM (Figure 4c). Western blot analysis further confirmed that MTD4-RBDV dose-dependently decreased the phosphorylation of key downstream signaling proteins, including Akt and MEK, with IC_50_ values of 1–3 μM (Figure 4d). Finally, flow cytometry analysis of H358 cells treated with MTD4-RBDV, followed by staining with AlexaFluor488-annexin V and propidium iodide, clearly indicated that cell death occurred via apoptosis (Figure S4). Collectively, these results establish MTD4-RBDV as a cell-permeable and biologically active Ras inhibitor.

**Figure 4.**
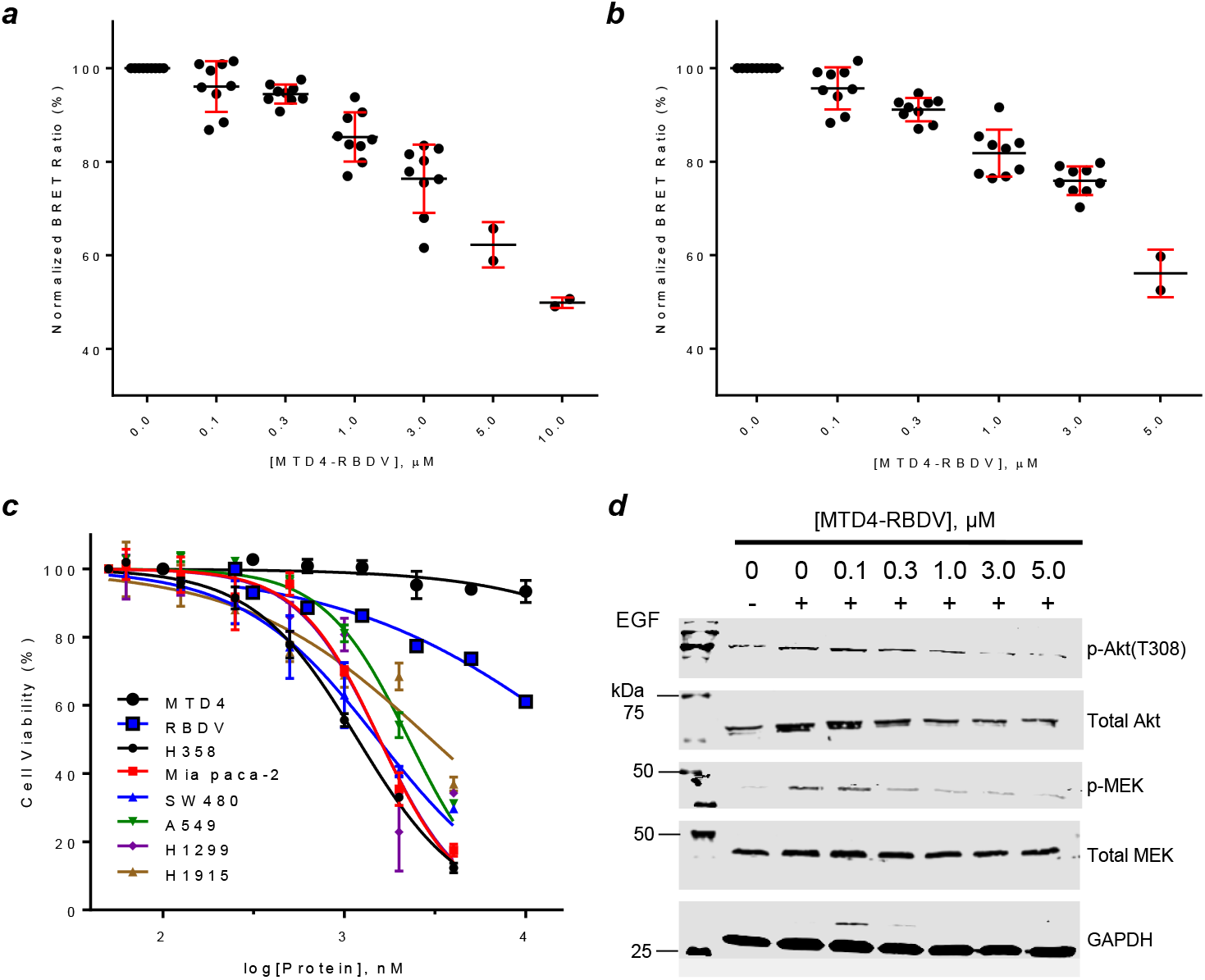
Inhibition of Ras signaling by MTD4-RBDV. (***a***) Dose-dependent reduction of the BRET signal in HEK293T cells transiently transfected with luciferase-KRasG12V and CRaf RBD-GFP by MTD4-RBDV (n = 9). (***b***) Same as (***a***), except cells were transfected with luciferase-KRasG12D. (***c***) Effect of MTD4-RBDV on the viability of the indicated cancer cell lines. MTD4 and RBDV controls were performed in H358 cells (n = 3). (***d***) Representative Western blots showing dose-dependent reduction of phosphorylated Akt and MEK in response to MTD4-RBDV.

### MTD4 Delivers Cargo Proteins in the Folded State

Intracellular delivery of cargo proteins in their native state is highly desirable. We investigated whether MTD4 delivers cargo proteins in the folded state, since some recent studies have reported that endosomal escape of certain CPPs (e.g., ZF5.3) requires the prior unfolding of both the CPP and its cargo.^30,31^ We generated a noncovalent complex of MTD4-HiBit and the large cargo protein LgBit-mCherry, through the high-affinity binding of HiBit to LgBit. Using live-cell confocal microscopy, we monitored the complex’s entry into HeLa cells. We stained both endosomes and lysosomes with specific markers to track their location.

As expected, cells treated with vehicle control (buffer only) showed no mCherry fluorescence (Figure 5). The HiBit/LgBit-mCherry complex on its own resulted in only a weak mCherry signal that largely colocalized with the endo/lysosomal compartments, indicating poor endosomal escape. In stark contrast, cells treated with the MTD4-HiBit/LgBit-mCherry complex showed intense, predominantly punctate mCherry fluorescence throughout the cytoplasm. Notably, only about 23% of the mCherry signal remained in the endo/lysosomal compartments, suggesting that the majority of the complex successfully reached the cytosol. The punctate mCherry fluorescence within the cytosol likely represents aggregates formed after the VBC-mediated escape, which may contain the protein complex along with membrane lipids and cellular proteins. The formation of these post-escape aggregates is a recently recognized bottleneck in the intracellular delivery of biomolecules.^43-45^

**Figure 5.**
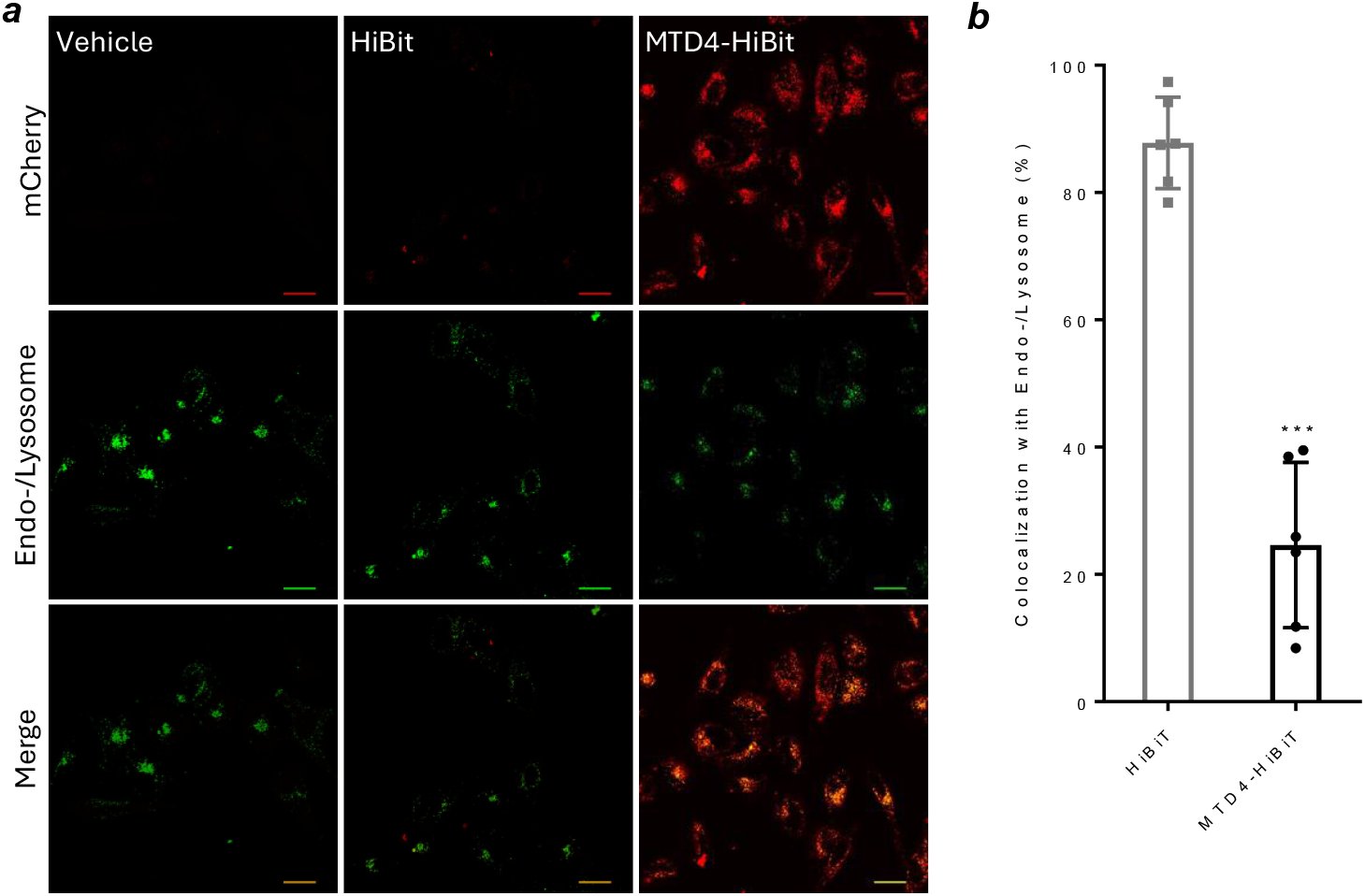
Intracellular delivery of LgBit-mCherry by MTD4-HiBit. (***a***) Representative live-cell confocal images of HeLa cells after treatment with vehicle (buffer only), 3 μM HiBiT + 4.5 μM LgBit-mCherry, or 3 μM MTD4–HiBiT + 4.5 μM LgBiT– mCherry. Prior to imaging, the cells were incubated with dextran–AlexaFluor647 and LysoTracker Deep Red, and their signals were acquired in the same channel and shown in green color. Scale bars, 20 μm. (***b***) Quantification of the fraction of mCherry signal colocalized with the Alexa647 channel (M2, shown as %). Data represent the mean ± SD of six independent experiments. ***, *p* < 0.001 (vs HiBiT).

Our finding that MTD4-HiBit successfully delivers LgBit-mCherry into the cell in a “piggyback” fashion confirms that all three components—MTD4, LgBit, and mCherry—remain folded during membrane translocation, as any unfolding would have caused LgBit-mCherry to dissociate from MTD4-HiBit. These results collectively demonstrate that MTD4-mediated delivery (presumably via the VBC mechanism) transports intact, folded proteins across the cell membrane.

### MTD4 Delivers Cre Recombinase into the Nucleus Ex Vivo and In Vivo

To evaluate MTD4’s capability for *in vivo* protein delivery and to gain insight into the biodistribution of MTD4 fusion proteins, we utilized a transgenic mouse model (Figure 6a). This system features a tdTomato gene preceded by stop codons flanked by two LoxP sites in its genome.^46^ We engineered an 80-kD fusion protein comprising MTD4, enhanced green fluorescent protein (EGFP), a Cre recombinase enzyme, and three nuclear localization sequences (NLS) [MTD4-EGFP-NLS-Cre-(NLS)_2_ or MEC]. Successful nuclear delivery of MEC into cells of these LoxP mice would result in the Cre-mediated excision of the stop codons, leading to the expression of tdTomato, a red fluorescent protein.

**Figure 6.**
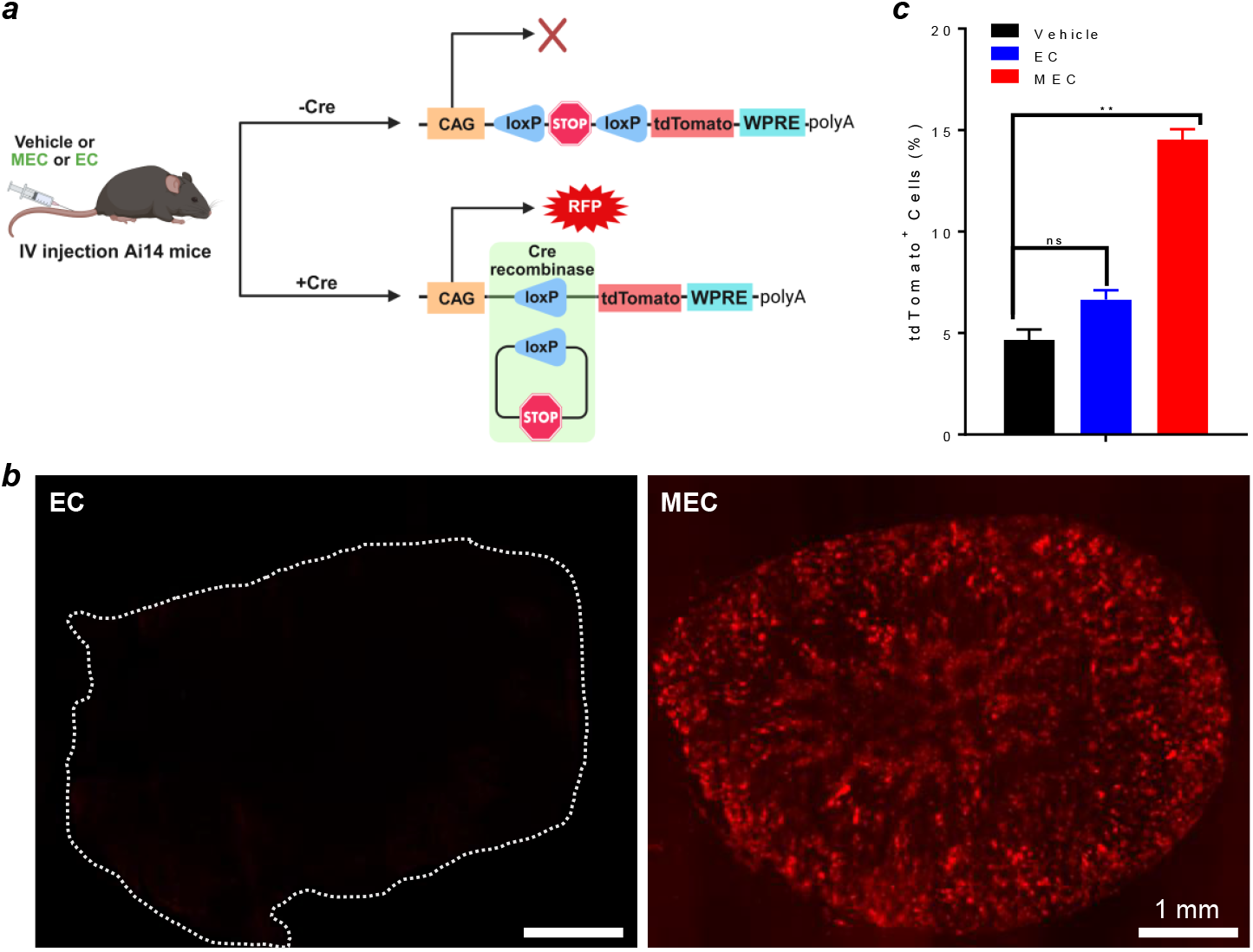
MEC-induced tdTomato expression in Ai14 mice. (***a***) Schematic representation of the Cre–loxP–tdTomato system in Ai14 mice. (***b***) Representative confocal microscopy images of cryosectioned kidneys (dotted line indicates tissue boundary) following intravenous injection of EC or MEC (98 µM, 100 µL). Scale bar: 1 mm. (***c***) Flow cytometry analysis of tdTomato-positive kidney cells from (***b***). Paired Student’s t-tests were performed between vehicle-, EC-, and MEC-treated groups (n = 3). **, p ≤ 0.01; ns, not significant (p > 0.05).

Initial ex vivo validation showed that treating primary cells isolated from LoxP mice with MEC resulted in robust tdTomato expression, whereas treatment with vehicle (buffer only) or a control protein lacking MTD4, EGFP-NLS-Cre-(NLS)_2_ (EC), did not. Flow cytometry analysis of these treated cells confirmed a dose-dependent expression of tdTomato, with ∼45% of cells becoming tdTomato positive at 1.5 μM MEC (Figure S5). Following this ex vivo success, LoxP mice were intravenously administered MEC (40 mg/kg), EC, or vehicle (buffer only) via tail vein injection and euthanized after 72 h. Tissues including the brain, heart, lung, kidney, liver, slow muscle (gastrocnemius), and fast muscle (soleus) were harvested, processed, and analyzed by confocal microscopy and flow cytometry. Robust tdTomato expression was observed in the kidneys (Figure 6b) and the injection site of MEC-treated mice, but not in other organs or in mice treated with vehicle or EC. Remarkably, the tdTomato fluorescence was distributed throughout the entire kidney organ. Flow cytometry analysis of cells derived from the homogenized kidney tissues revealed that ∼15% of all kidney cells of MEC-treated mice were tdTomato positive, as opposed to ∼5% for EC- or vehicle-treated mice (Figure 6c). These results demonstrate the *in vivo* delivery potential of MTD4 fusion proteins. The lack of significant tdTomato expression in other tissues was likely due to the rapid proteolytic degradation of MEC *in vivo*, resulting in limited exposure.

### MTD4 Enables Intracellular Delivery and Functional Replacement of ASL

To test MTD4’s potential for delivering therapeutic proteins, we engineered ASL-MTD4, a fusion protein consisting of human argininosuccinate lyase (ASL) fused at its N-terminus to MTD4. This construct serves as a candidate enzyme replacement therapy for ASL deficiency (ASLD), a urea cycle disorder characterized by elevated levels of argininosuccinate and hyperammonemia.^47^ ASL is a cytosolic enzyme crucial for the fourth step of the urea cycle, catalyzing the conversion of argininosuccinate into arginine and fumarate. We first assessed the cellular uptake of ASL-MTD4 in ASL-deficient primary human fibroblasts (GM00525). Western blot analysis revealed the efficient uptake of ASL-MTD4 at 500 nM, while ASL (no MTD4) showed no detectable internalization (Figure 7a). To confirm in-cellulo enzymatic activity, we measured the intracellular argininosuccinate levels in GM00525 cells post-treatment. ASL-MTD4 dose-dependently reduced the argininosuccinate concentration to levels comparable to those found in wild-type primary human fibroblasts (GM01661), confirming the functional delivery of catalytically active ASL (Figure 7b).

**Figure 7.**
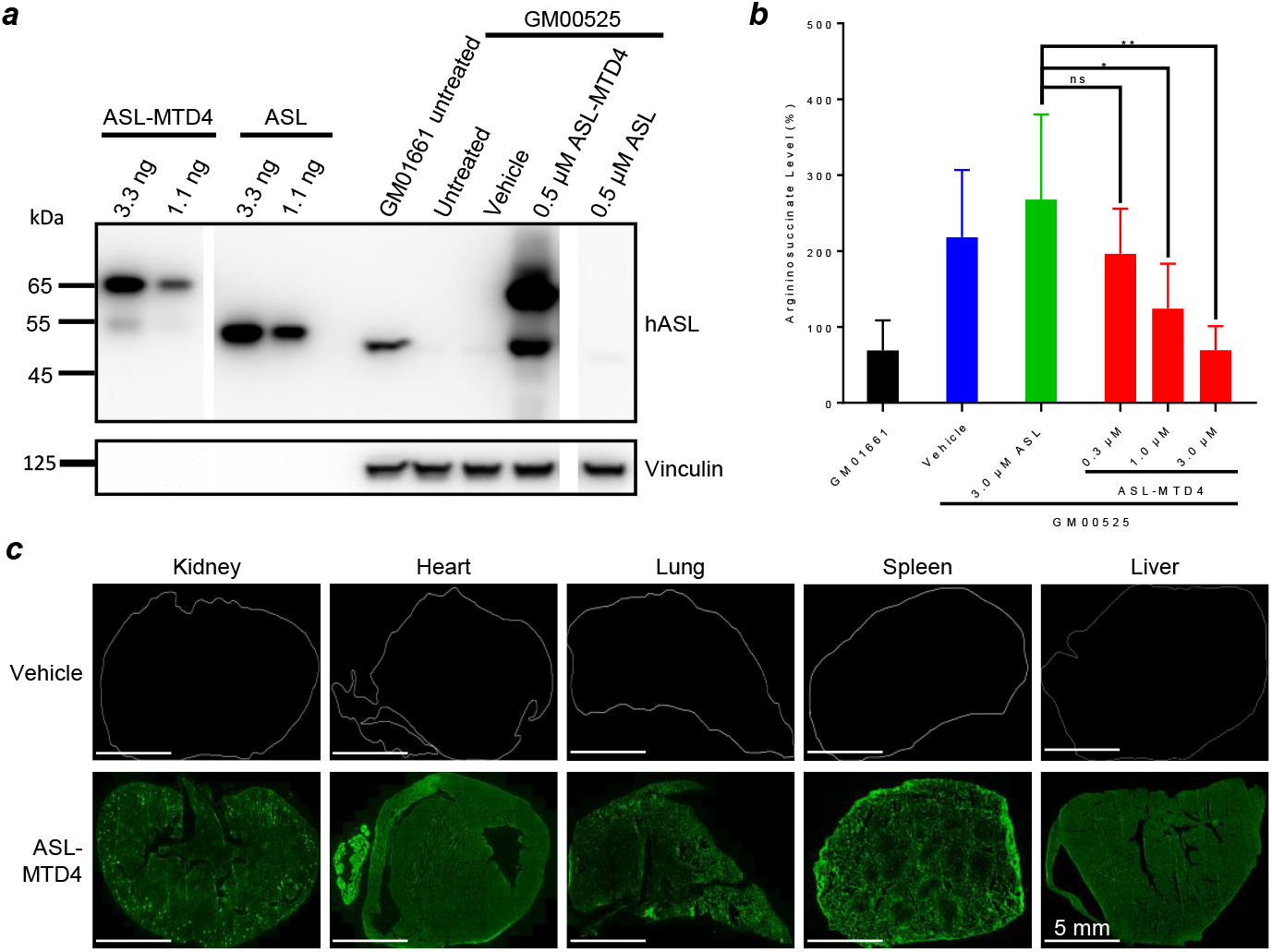
ASL-MTD4 exhibits cytosolic uptake ex vivo and broad biodistribution in mice. (***a***) Representative Western blot analysis of ASL and ASL-MTD4 in healthy human primary fibroblasts (GM01661) and ASLD patient-derived fibroblasts (GM00525) after treatment with null, vehicle (buffer only), ASL, or ASL-MTD4. The lower band in ASL-MTD4 lanes corresponds to a proteolytic fragment. (***b***) Quantification of intracellular argininosuccinate levels in untreated GM01661 cells or GM00525 cells treated with vehicle, ASL (3.0 µM), or ASL-MTD4 (0.3, 1.0, or 3.0 µM). Error bars represent mean ± SD of six independent experiments (biological replicates). *p ≤ 0.05; **p ≤ 0.01; ns, not significant (p > 0.05). (***c***) Representative confocal microscopy images of cryosectioned mouse organs collected 4 h after intravenous administration of ASL-MTD4 (800 µg) or vehicle. Sections were immunostained with anti-human ASL antibody (PA5-117795), with green fluorescence indicating ASL-MTD4 localization. Scale bars: 5 mm.

For *in vivo* assessment, mice were intravenously injected with equimolar amounts of ASL-MTD4 or ASL. ASL-MTD4 cleared rapidly from the serum, becoming undetectable ∼3 h post-injection, but was readily detectable in multiple organs at 4 h. In stark contrast, ASL remained predominantly in circulation with negligible tissue uptake. Confocal microscopy of liver and kidney sections corroborated the uptake of ASL-MTD4, but not ASL, into these tissues (Figure S6a). Finally, to investigate the broader biodistribution, mice were intravenously injected with 40 mg/kg of ASL-MTD4, and major organs were collected 4 h post-injection. Confocal imaging of tissue sections revealed broad biodistribution and remarkably uniform cellular uptake across the liver, kidney, lung, heart, and spleen (Figure 7c). High-magnification images within these tissues consistently showed a diffused ASL signal within most cells, strongly indicating successful cytosolic delivery (Figure S6b).

### Advantages and Disadvantages of MTDs

Several key attributes render the MTD platform ideally suited for the intracellular delivery of peptides and proteins, both *in vitro* and *in vivo*. First, the MTDs are exceptionally versatile; essentially any peptide or protein may, in principle, be rendered cell permeable by genetically fusing an MTD to its N-or C-terminus. The versatility is also reflected by the fact that the MTDs can penetrate virtually any eukaryotic cell that undergoes active endocytosis, including mammalian, plant, and fungal cells, although for plant and fungal cells the size of the cargo is generally limited to ≤40 kDa due to the presence of a cell wall. To date, we have used MTDs to deliver over 30 different peptides and proteins, ranging from 0.4 to 260 kDa in molecular weight and 5.7 to 11 in isoelectric point (pI), into mammalian, plant, or fungal cells. Our data indicates that fusion with MTDs generally does not adversely affect the function or activity of a protein. Second, MTD4 is highly active, demonstrating a cytosolic delivery efficiency rivaling or even surpassing that of CPP12,^23^ a potent cyclic CPP previously developed in one of our laboratories which is currently in clinical development. Notably, MTD4 remains very active at low concentrations, making it particularly useful for delivering highly potent proteins such as gene-editing nucleases and other therapeutic enzymes. Third, the MTDs were designed to bind directly to membrane phospholipids (as opposed to protein receptors), enter the cell by endocytosis, and subsequently escape the endosome by the vesicle budding-and-collapse (VBC) mechanism.^35-37^ Since all cells share similar membrane phospholipids, this design strategy enables the MTDs to access a broad range of tissues and cell types following systemic administration. In contrast, leading delivery technologies such as lipid nanoparticles and viral vectors are largely limited to the liver. Remarkably, MTD4 fusion proteins demonstrated deep tissue penetration, resulting in a homogeneous distribution throughout the organs, which is rarely seen with other delivery technologies. Fourth, the MTD4 core domain (MTD4s) is thermodynamically (T_M_ = 53 ± 2 °C) as well as proteolytically stable (t_1_/_2_ > 24 h in human serum) and can be readily expressed in *Escherichia coli* with high yields. These properties greatly facilitate the straightforward and scalable production of MTD4 fusion proteins for clinical and other applications. Lastly, MTD4 is minimally engineered from a widely expressed, extracellular human matrix protein domain (FN3),^33^ involving only six-amino acid substitutions across two loops. Hence, MTD4 is expected to have low immunogenicity, which is corroborated by our in-silico analysis (Figure S7) and the generally low immunogenicity observed for other FN3-based therapeutics.^48^

The MTD platform has some limitations. While MTD4s is proteolytically stable, it does not protect cargo proteins from proteolytic degradation, unlike encapsulation-based delivery platforms (e.g., LNPs). Thus, MTDs are practically limited to delivering proteins of sufficient thermodynamic and proteolytic stabilities. A case in point is MEC, which undergoes rapid proteolysis in human serum, most likely within the internal NLS region. As such, while MEC resulted in robust tdTomato expression in mouse primary cells ex vivo at a concentration of 150 nM (Fig. S5), it failed to achieve significant *in vivo* gene editing in most mouse tissues outside the kidney and the injection site, even at a 40 mg/kg dose. In contrast, the relatively stable ASL-MTD4 tetramer (∼260 kDa) exhibited broad biodistribution (Fig. 7c). Potential strategies to mitigate these limitations include implementing PEGylation^49^ or PASylation^50^ to improve stability and reduce renal clearance, particularly since MTD4’s delivery abilities are largely independent of the cargo’s biophysical properties. We also observed that certain MTD-cargo combinations may result in reduced or abolished uptake, possibly due to intramolecular interactions between the CPP loops of MTD and the cargo protein. These effects might be mitigated by incorporating alternative linker sequences, altering the relative orientation (i.e., switching MTD from the N- to C-terminus of cargo or vice versa), or inserting an intervening domain between MTD and cargo. Predictive protein modeling tools such as Phyre2^51^ or AlphaFold^52^ may be leveraged to identify and circumvent such incompatibilities. Ongoing studies in our laboratories are addressing these limitations, with the goal of refining the MTD platform into a general delivery vehicle for peptides, proteins, and other biomolecules (e.g., siRNAs).

## CONCLUSION

In this work, we have developed a novel delivery platform, MTDs, and demonstrated its utility for the intracellular delivery of small molecules (e.g., TMR), peptides (e.g., HiBit), and proteins. MTD4 can be recombinantly fused to the N- or C-terminus of any peptide or protein and produced in *E. coli* or other hosts with high yields. Its high cytosolic delivery efficiency across a broad range of concentrations and excellent proteolytic stability make it well suited for the intracellular delivery of peptides, proteins, and potentially other biomolecules as therapeutics as well as biomedical research tools.

## ASSOCIATED CONTENT

### Supporting Information

Experimental details, Figures S1–S7, Table S1,

References. This material is available free of charge via the Internet at http://pubs.acs.org.

## AUTHOR INFORMATION

### Corresponding Author

*Dehua Pei-* Department of Chemistry and Biochemistry, The Ohio State University, 100 West 18th Avenue, Columbus, 43210, Ohio, United States

### Author Contributions

D.P. conceived the project and designed the MTD platform. D.P. and P.B. designed most experiments. J.G., P.B., V.L.K., and L.T. designed the ASL study; P.B., D.P., J.Z., N.L., and Q.E.W. designed the Cre–loxP study. J.L.R., H.S., P.B., B.H., and T.P. cloned plasmids and expressed, purified, or synthesized proteins and peptides. H.S. performed CellTiter-Glo assays, apoptosis assays, and Western blotting for the RBD constructs. P.B. performed CellTiter-Glo assays, Western blotting, HiBiT assays, flow cytometry, and confocal microscopy for all other experiments, except for the ASL study, which was conducted by WuXi AppTec (China). N.L. performed organ harvesting and *in vivo* injections.D.P. and P.B. analyzed all data, with contributions from J.G. and V.L.K. (ASL study) and Q.E.W. and N.L. (Cre–loxP study). D.P. and P.B. wrote the manuscript.

## ACKNOWLEDGMENT

We acknowledge resources from the Campus Microscopy and Imaging Facility (CMIF) and the OSU Comprehensive Cancer Center (OSUCCC) Microscopy Shared Resource (MSR), The Ohio State University (RRID:SCR_025078). This facility is supported in part by grant P30 CA016058, National Cancer Institute, Bethesda, MD. We thank the Comparative Pathology & Digital Imaging Shared Resource (CPDISR) at The Ohio State University Comprehensive Cancer Center, Columbus, OH for whole slide imaging, the research at CPDISR is supported by The Ohio State University Comprehensive Cancer Center and the National Institutes of Health under grant number P30 CA016058. We thank the Flow Cytometry Shared Resource (FCSR), and the University Laboratory Animal Resources (ULAR) at The Ohio State University for their technical support. This work was supported by the National Institutes of Health (GM122459).

## AUTHOR DECLARATIONS

D.P. and P.B. are co-inventors on patent applications related to the MTD platform filed by The Ohio State University. D.P. is cofounder, consultant, and equity owner of Permeasis Therapeutics and Scioto AgriTech, which have licensed MTD technology from OSU. P.B. is a consultant, and L.T., J.G., and V.L.K. are employees of Permeasis Therapeutics. The other authors declare no competing financial interest.

---

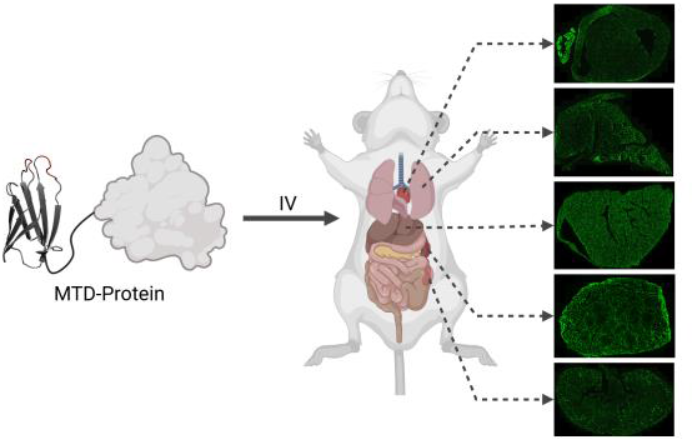

## REFERENCES

1. Silva, A. K. A.; Sagné, C.; Gazeau, F.; Abasolo, I. Enzyme Replacement Therapy: Current Challenges and Drug Delivery Prospects via Extracellular Vesicles. Rare Dis. Orphan Drugs J. 2022, 1, 13.

2. van der Oost, J.; Constantinos, P. The Genome Editing Revolution. Trends Biotechnol. 2023, 41, 396–409.

3. Higueruelo, A. P.; Jubb, H.; Blundell, T. L. Protein–Protein Interactions as Druggable Targets: Recent Technological Advances. Curr. Opin. Pharmacol. 2013, 13, 791–796.

4. VanDyke, D.; Taylor, J. D.; Kaeo, K. J.; Hunt, J.; Spangler, J. B. Biologics-Based Degraders—An Expanding Toolkit for Targeted Protein Degradation. Curr. Opin. Biotechnol. 2022, 78, 102807.

5. Khan, M. M.; Filipczak, N.; Torchilin, V. P. Cell-Penetrating Peptides: A Versatile Vector for Co-Delivery of Drugs and Genes in Cancer. J. Control. Release 2021, 330, 1220–1228.

6. Beilhartz, G. L.; Sugiman-Marangos, S. N.; Melnyk, R. A. Repurposing Bacterial Toxins for Intracellular Delivery of Therapeutic Proteins. Biochem. Pharmacol. 2017, 142, 13–20.

7. Zhao, Z.; Anselmo, A. C.; Mitragotri, S. Viral Vector-Based Gene Therapies in the Clinic. Bioeng. Transl. Med. 2022, 7, e10258.

8. Ita, K. Polyplexes for Gene and Nucleic Acid Delivery: Progress and Bottlenecks. Eur. J. Pharm. Sci. 2020, 150, 105358.

9. Filipczak, N.; Pan, J.; Yalamarty, S. S. K.; Torchilin, V. P. Recent Advancements in Liposome Technology. Adv. Drug Deliv. Rev. 2020, 156, 4–22.

10. Mitchell, M. J.; Billingsley, M. M.; Haley, R. M.; Wechsler, M. E.; Peppas, N. A.; Langer, R. Engineering Precision Nanoparticles for Drug Delivery. Nat. Rev. Drug Discov. 2021, 20, 101–124.

11. Wang, Y. S.; Zhang, Z.; Luo, J.; et al. mRNA-Based Vaccines and Therapeutics: An In-Depth Survey of Current and Upcoming Clinical Applications. J. Biomed. Sci. 2023, 30, 84.

12. Wang, D.; Tai, P. W. L.; Gao, G. Adeno-Associated Virus Vector as a Platform for Gene Therapy Delivery. Nat. Rev. Drug Discov. 2019, 18, 358–378.

13. Khoshnood, S.; et al. Bacteria-Derived Chimeric Toxins as Potential Anticancer Agents. Front. Oncol. 2022, 12, 953678.

14. Kumar, M.; Kulkarni, P.; Liu, S.; Chemuturi, N.; Shah, D. K. Nanoparticle Biodistribution Coefficients: A Quantitative Approach for Understanding the Tissue Distribution of Nanoparticles. Adv. Drug Deliv. Rev. 2023, 194, 114708.

15. Dhungel, B. P.; et al. Understanding AAV Vector Immunogenicity: From Particle to Patient. Theranostics 2024, 14, 1260–1288.

16. Pei, D.; Buyanova, M. Overcoming Endosomal Entrapment in Drug Delivery. Bioconjug. Chem. 2019, 30, 273–283.

17. Fawell, S.; et al. Tat-Mediated Delivery of Heterologous Proteins into Cells. Proc. Natl. Acad. Sci. U. S. A. 1994, 91, 664–668.

18. Patel, S. G.; et al. Cell-Penetrating Peptide Sequence and Modification Dependent Uptake and Subcellular Distribution of Green Fluorescent Protein in Different Cell Lines. Sci. Rep. 2019, 9, 6298.

19. Ando, Y.; Nakazawa, H.; Miura, D.; Otake, M.; Umetsu, M. Enzymatic Ligation of an Antibody and Arginine 9 Peptide for Efficient and Cell-Specific siRNA Delivery. Sci. Rep. 2021, 11, 21882.

20. Guidotti, G.; et al. Cell-Penetrating Peptides: From Basic Research to Clinics. Trends Pharmacol. Sci. 2017, 38, 406–424.

21. Lättig-Tünnemann, G.; et al. Backbone Rigidity and Static Presentation of Guanidinium Groups Increases Cellular Uptake of Arginine-Rich Cell-Penetrating Peptides. Nat. Commun. 2011, 2, 453.

22. Mandal, D.; Shirazi, S. N.; Parang, K. Cell-Penetrating Homochiral Cyclic Peptides as Nuclear-Targeting Molecular Transporters. Angew. Chem., Int. Ed. 2011, 50, 9633–9637.

23. Qian, Z.; et al. Discovery and Mechanism of Highly Efficient Cyclic Cell-Penetrating Peptides. Biochemistry 2016, 55, 2601–2612.

24. Zhang, W.; Lin, M.; Yan, Q.; et al. An Intracellular Nanobody Targeting T4SS Effector Inhibits Ehrlichia Infection. Proc. Natl. Acad. Sci. U. S. A. 2021, 118, e2024102118.

25. Li, X.; Kheirabadi, M.; Dougherty, P. G.; et al. The Endosomal Escape Vehicle Platform Enhances Delivery of Oligonucleotides in Preclinical Models of Neuromuscular Disorders. Mol. Ther. Nucleic Acids 2023, 33, 273–285.

26. Spicer, C. D.; Davis, B. G. Selective Chemical Protein Modification. Nat. Commun. 2014, 5, 4740.

27. Appelbaum, J. S.; LaRochelle, J. R.; Smith, B. A.; Balkin, D. M.; Holub, J. M.; Schepartz, A. Arginine Topology Controls Escape of Minimally Cationic Proteins from Early Endosomes to the Cytoplasm. Chem. Biol. 2012, 19, 819–830.

28. Shen, F.; Zheng, G.; Setegne, M.; Tenglin, K.; Izadi, M.; Xie, H.; Zhai, L.; Orkin, S. H.; Dassama, L. M. K. A Cell-Permeant Nano-body-Based Degrader That Induces Fetal Hemoglobin. ACS Cent. Sci. 2022, 8, 1695–1703.

29. Zhang, X.; Cattoglio, C.; Zoltek, M.; Vetralla, C.; Mozumdar, D.; Schepartz, A. Dose-Dependent Nuclear Delivery and Transcriptional Repression with a Cell-Penetrant MeCP2. ACS Cent. Sci. 2023, 9, 277–288.

30. Zoltek, M.; Vázquez Maldonado, A. L.; Zhang, X.; Dadina, N.; Lesiak, L.; Schepartz, A. HOPS-Dependent Endosomal Escape Demands Protein Unfolding. ACS Cent. Sci. 2024, 10, 860–870.

31. Giudice, J.; Brauer, D. D.; Zoltek, M.; Vázquez-Maldonado, A. L.; Dadina, N.; Kelly, M.; Schepartz, A. The Biophysical Requirements That Govern the Efficient Endosomal Escape of Designed Mini-Proteins. Nat. Chem. 2025, 17, 1227–1235.

32. Chen, K.; Pei, D. Engineering Cell-Permeable Proteins through Insertion of Cell-Penetrating Motifs into Surface Loops. ACS Chem. Biol. 2020, 15, 2568–2576.

33. Koide, A.; Bailey, C. W.; Huang, X.; Koide, S. The Fibronectin Type III Domain as a Scaffold for Novel Binding Proteins. J. Mol. Biol. 1998, 284, 1141–1151.

34. Koide, S.; Koide, A.; Lipovšek, D. Target-Binding Proteins Based on the 10th Human Fibronectin Type III Domain (10Fn3). Methods Enzymol. 2012, 503, 135–156.

35. Sahni, A.; Qian, Z.; Pei, D. Cell-penetrating Peptides Escape the Endosome by Inducing Vesicle Budding and Collapse. ACS Chem. Biol. 2020, 15, 2485–2492.

36. Sahni, A.; Ritchey, J. L.; Qian, Z.; Pei, D. Cell-Penetrating Peptides Translocate across the Plasma Membrane by Inducing Vesicle Budding and Collapse. J. Am. Chem. Soc. 2024, 146, 25371–25382.

37. Pei, D. How Do Biomolecules Cross the Cell Membrane? Acc. Chem. Res. 2022, 55, 309–318.

38. Kim, J. S.; et al. Endosomal Acidic pH-Induced Conformational Changes of a Cytosol-Penetrating Antibody Mediate Endosomal Escape. J. Control. Release 2016, 235, 165–175.

39. Schwinn, M. K.; Machleidt, T.; Zimmerman, K.; et al. CRISPR-Mediated Tagging of Endogenous Proteins with a Luminescent Peptide. ACS Chem. Biol. 2018, 13, 467–474.

40. Tonks, N. K. PTP1B: From the Sidelines to the Front Lines! FEBS Lett. 2003, 546, 140–148.

41. Wiechmann, S.; et al. Conformation-Specific Inhibitors of Activated Ras GTPases Reveal Limited Ras Dependency of Patient-Derived Cancer Organoids. J. Biol. Chem. 2020, 295, 4526–4540.

42. Bery, N.; et al. BRET-Based RAS Biosensors Show That a Novel Small Molecule Inhibits RAS–Effector Protein–Protein Interactions. eLife 2018, 7, e37122.

43. Pei, D. Endosomal Escape of Lipid Nanoparticles: A New Perspective on the Literature Data. ChemRxiv 2025, DOI 10.26434/chemrxiv-2025-t8115.

44. Cheung, T H.; Shoichet, M. S. The Interplay of Endosomal Escape and RNA Release from Polymeric Nanoparticles. Langmuir 2025, 41, 7174–7190.

45. Liu, H.; Chen, M. Z.; Payne, T.; Porter, C. J. H.; Pouton, C. W.; Johnston, A. P. R. Beyond the Endosomal Bottleneck: Understanding the Efficiency of mRNA/LNP Delivery. Adv. Funct. Mater. 2024, 34, 2404510.

46. Madisen, L.; Zwingman, T. A.; Sunkin, S. M.; et al. A Robust and High-Throughput Cre Reporting and Characterization System for the Whole Mouse Brain. Nat. Neurosci. 2010, 13, 133–140.

47. Jalil, S.; Keskinen, T.; Juutila, J.; et al. Genetic and Functional Correction of Argininosuccinate Lyase Deficiency Using CRISPR Adenine Base Editors. Am. J. Hum. Genet. 2024, 111, 714–728.

48. Iwamoto, N.; Sato, Y.; Manabe, A.; Inuki, S.; Ohno, H.; Nonaka, M.; Oishi, S. Design and Synthesis of Monobody Variants with Low Immunogenicity. ACS Med. Chem. Lett. 2023, 14, 1596–1601.

49. Veronese, F. M. Peptide and Protein PEGylation: A Review of Problems and Solutions. Biomaterials 2001, 22, 405–417.

50. Gebauer, M.; et al. PASylation: A Biological Alternative to PEGylation for Extending the Plasma Half-Life of Pharmaceutically Active Proteins. Protein Eng. Des. Sel. 2013, 26, 489–501.

51. Kelley, L. A.; et al. The Phyre2 Web Portal for Protein Modeling, Prediction and Analysis. Nat. Protoc. 2015, 10, 845–858.

52. Jumper, J.; et al. Highly Accurate Protein Structure Prediction with AlphaFold. Nature 2021, 596, 583–589.

